# Individual differences in developmental trajectory leave a male polyphenic signature in bulb mite populations

**DOI:** 10.1101/2023.02.06.527265

**Authors:** Jacques A. Deere, Isabel M. Smallegange

**Affiliations:** Institute for Biodiversity and Ecosystem Dynamics, University of Amsterdam, PO Box 94240, 1090 GE Amsterdam, the Netherlands; School of Natural and Environmental Sciences, Newcastle University, Newcastle upon Tyne, NE1 7RU, UK

**Keywords:** alternative male phenotypes, dispersal, eco-evolutionary dynamics, male morph coexistence, polyphenism

## Abstract

Developmental plasticity alters phenotypes and can in that way change the response to selection. When alternative phenotypes show different life history trajectories, developmental plasticity can also affect, and be affected by, population size-structure in an eco-evolutionary interaction. Developmental plasticity often functions to anticipate future conditions but it can also mitigate current stress conditions. Both types of developmental plasticity have evolved under different selections and this raises the question if they underlie different eco-evolutionary population dynamics. Here, we tested, in a long-term population experiment using the male polyphenic bulb mite (*Rhizoglyphus robini*), if the selective harvesting of juveniles of different developmental stages concurrently alters population size (ecological response) and male adult phenotype expression (evolutionary response) in line with eco-evolutionary predictions that assume the male polyphenism is anticipatory or mitigating. We found that the frequency of adult males that expressed costly (fighter) morphology was lowest under the most severe juvenile harvesting conditions. This response cannot be explained if we assume that adult male phenotype expression is to anticipate adult (mating) conditions because, in that case, only the manipulation of adult performance would have an effect. Instead, we suggest that juveniles mitigate their increased mortality risk by expediating ontogeny to forego the development of costly morphology and mature quicker but as a defenceless scrambler. If, like in mammals and birds where early-life stress effects are extensively studied, we account for such pre-adult viability selection in coldblooded species, it would allow us to (i) better characterise natural selection on trait development like male polyphenisms, (ii) understand how it can affect the response to other selections in adulthood, and (iii) understand how such trait dynamics influence, and are influenced by, population dynamics.

## Introduction

Across the animal kingdom, developmental plasticity comprises a switch to alternative developmental pathways that depends on an individual’s condition – the resource budget available for the production and maintenance of adaptive traits (Hill 2011; Casasa et al. 2020; Nijhout & McKenna 2018) – and which can lead to food-dependent allometric plasticity and the adaptive expression of alternative phenotypes in adulthood (polyphenism) (West-Eberhard 2003; Nijhout & McKenna 2018). For example, many intraspecific, arthropod male dimorphisms comprise large majors that have weapons for male-male competition, and smaller, weaponless minors that sneak matings (Moczek 2003; Smallegange 2011; Buzatto & Machado 2014). Importantly, these adaptive, condition-dependent alternative phenotypes typically have different survival, growth and reproduction rates because they follow different developmental trajectories (West-Eberhard 2003; Smallegange 2011; Weir et al. 2016). Differences in individual development, be they plastic or genetic, will thus affect a population’s age or life stage structure, size and growth. Such population changes can in turn influence individual development through processes like density-dependent competition over resources (food) and density-dependent selection (Travis et al. 2013), creating eco-evolutionary feedbacks at the population level (Smallegange & Deere 2014; Croll et al. 2019; Smallegange 2022). However, accurately predicting the population dynamical responses to perturbation of developmental trajectories is still difficult (Smallegange et al. 2019; Smallegange 2022).

The way in which a perturbation to individual development can impact the eco-evolutionary dynamics of populations depends on what selection pressures are assumed to act on adaptive developmental plasticity (Smallegange 2022). Different selection pressures determine different types of adaptive developmental plasticity (e.g., Forsman 2015; Nettle and Bateson 2015) and here we focus on two distinct types: anticipatory and mitigating adaptive developmental plasticity (Smallegange et al. 2019; Smallegange 2022). Specifically, in case of the minor and major males described above, whether a male develops into an armed major male or an unarmed minor male depends on whether it reaches a critical resource budget threshold, assumed to be approximated by a body size threshold, during development (Plaistow et al. 2005; Sasson et al. 2016; Cotton et al. 2004). If the adaptive developmental plasticity is *anticipatory*, this condition-cued threshold mechanism can be interpreted as the evolutionary result of disruptive sexual selection on alternative mating tactics (West-Eberhard 2003; Smallegange et al. 2019): good-condition juvenile males maximise fitness by becoming majors, anticipating competitive mating success; bad-condition juveniles anticipate failure at competitive success and salvage fitness by becoming minors (Shuster & Wade 2003). On the other hand, the condition-cued threshold mechanism could be part of a heritable developmental system that adaptively mitigates stressful environmental conditions (Badyaev 2005, Del Giudice et al. 2018; Ellis & Del Giudice 2019). Such *mitigating* adaptive developmental plasticity is the result of natural selection that favours developmental systems that tend to construct phenotypes that are successful, relative to other variant systems, at surviving and reproducing (Griffiths & Gray 1994). This means that juvenile males can mitigate stress during maturation by refraining from developing costly morphology to maintain fitness and mature and survive as an adult minor, as opposed to risk dying during maturation when investing in costly major morphology (Smallegange et al. 2019). To summarise, in both kinds of adaptive plasticity, the fitness functions of the alternative male phenotypes cross over a condition gradient, but anticipatory adaptive developmental plasticity assumes that the threshold for alternative male phenotype expression at the cross-section of the fitness functions is evolutionarily regulated by how adult males perform, whereas mitigating adaptive developmental plasticity assumes that the threshold is regulated by the dynamics of population density, food competition and individual resource budgets.

The fitness functions of both types of developmental plasticity are formalised in the environmental threshold (ET) model (Hazel et al. 1990; 2004). Under the ET model, the alternative expression of phenotypes has been hypothesized to be an environmentally cued threshold trait. It is assumed that threshold traits are based on a continuously distributed, polygenic trait, called the ‘liability’ and a threshold of expression such that individuals that are above this threshold express one phenotype while those below the threshold express the alternative (Roff 1996). Candidate traits for the liability are, for example, hormone profiles (Roff 1996). The position of the threshold depends on a cue that reliably correlates with the status of the environment, which in many taxa represents condition (Tomkins et al. 2011; Smallegange & Deere 2014). The ET model assumes that, in response to environment-specific selection, alternative phenotypes have evolved different fitness functions, through which selection can affect the distribution of individual liabilities. Because alternative phenotype frequency depends on the distribution of individual liabilities and the cue distribution, both are taken into account in determining how phenotype fitness influences the evolution of liabilities and hence the evolution of a threshold trait. The threshold thus evolves if the fitness functions change in response to changing selection(s), evolutionary impacting alternative phenotype expression. Furthermore, the cue (here: condition) distribution within the ET model informs on population size and structure (Smallegange & Coulson 2013). If the cue (here: condition) distribution, and thus population size and structure, changes, it ecologically impacts what proportion of individuals expresses which alternative phenotype (e.g., a reduction in overall condition reduces the frequency of males that can reach a high enough condition to develop into a fighter). Thus, the ET model gives us the ingredients to create predictions on how changes to the fitness functions and condition distribution of anticipatory or mitigating developmental plasticity fuel eco-evolutionary population change.

We conducted a long-term population experiment to test hypotheses on how the selective removal of juveniles of different condition in populations of the male polyphenic bulb mite *Rhizoglyphus robini* impacts eco-evolutionary population responses through anticipatory and mitigating developmental plasticity, under the assumptions of the ET model. Adult fighter males of the bulb mite possess a proportionally thickened third leg pair with dagger-like claws that are used in fights to kill rival males (Radwan et al. 2000). Whereas fighter males metamorphose from good-condition juveniles that have a large resource budget, juveniles that are in a bad condition (i.e., have a small resource budget) when metamorphosing into an adult instead express the scrambler phenotype that does not have such leg modifications (Rhebergen et al. 2022). Note that male adult phenotype expression does not depend on pheromone cues that inform on the frequency of each male phenotype in the population (Deere & Smallegange 2014) or mite density (Radwan 1995).

Further, in response to unfavourable environmental conditions (*e.g.*, low food), *R. robini* expresses a facultative dispersal morph during development, called the deutonymph, which does not feed and is adapted to phoretic dispersal (Díaz et al. 2000). Deutonymph expression occurs mid-development, prior to the final instar stage from which juveniles metamorphose into adults (Díaz et al. 2000), and, thus, prior to when adult male phenotype is determined (which is immediately before metamorphosis [Rhebergen et al. 2022]). Because deutonymph expression carries developmental costs, including reduced size at metamorphosis in both sexes and reduced egg production in females (Deere et al. 2015), our working assumption is that only individuals that are of sufficiently good condition can develop into one if cued by poor environmental conditions. This assumption is supported by the fact that in two of our previous studies (Smallegange & Coulson 2011; Deere et al. 2015), male deutonymphs always matured as a fighter, from which we surmise that males always were in sufficiently good condition to develop costly fighter legs.

In our experiment, we selectively removed individuals of different conditions by removing deutonymphs, or other juveniles. From the ET model, assuming that the male polyphenism is anticipatory, we predict that the ecological response of removing deutonymphs from the population is that it will lower the righthand side of the distribution of individual conditions (because individuals of relatively good condition are removed), and thus population size (compare blue dotted distribution size [before harvesting] with red dashed distribution size [after harvesting for several generations] in Fig. 1c). However, we expect that this will not elicit an evolutionary response because male fitness is assumed to be unaffected by juvenile performance (it is only affected by adult male performance). Thus, we expect that the mean threshold for fighter expression, and thus the proportion of males that are fighters, will track the latter change in the condition distribution because relative, and not absolute condition is important in determining alternative phenotype fitness (Tomkins et al. 2011) (compare the proportions of the differently shaded areas under the condition distribution in Fig. 1a [before harvesting] with Fig. 1c [after harvesting for several generations]). Likewise, we predict that the selective removal of juveniles (bar deutonymphs) of random condition has no effect on fighter expression (compare the proportions of the differently shaded areas under the condition distribution in Fig. 1a [before harvesting] with Fig. 1e [after harvesting for several generations]), because fitness functions remain unaltered. The reduction in population size because of the harvesting of juveniles, however, will be reflected in an overall reduced size of the condition distribution (compare blue dotted distribution size [before harvesting] with red dashed distribution size [after harvesting for several generations] in Fig. 1e).

**Figure 1.**
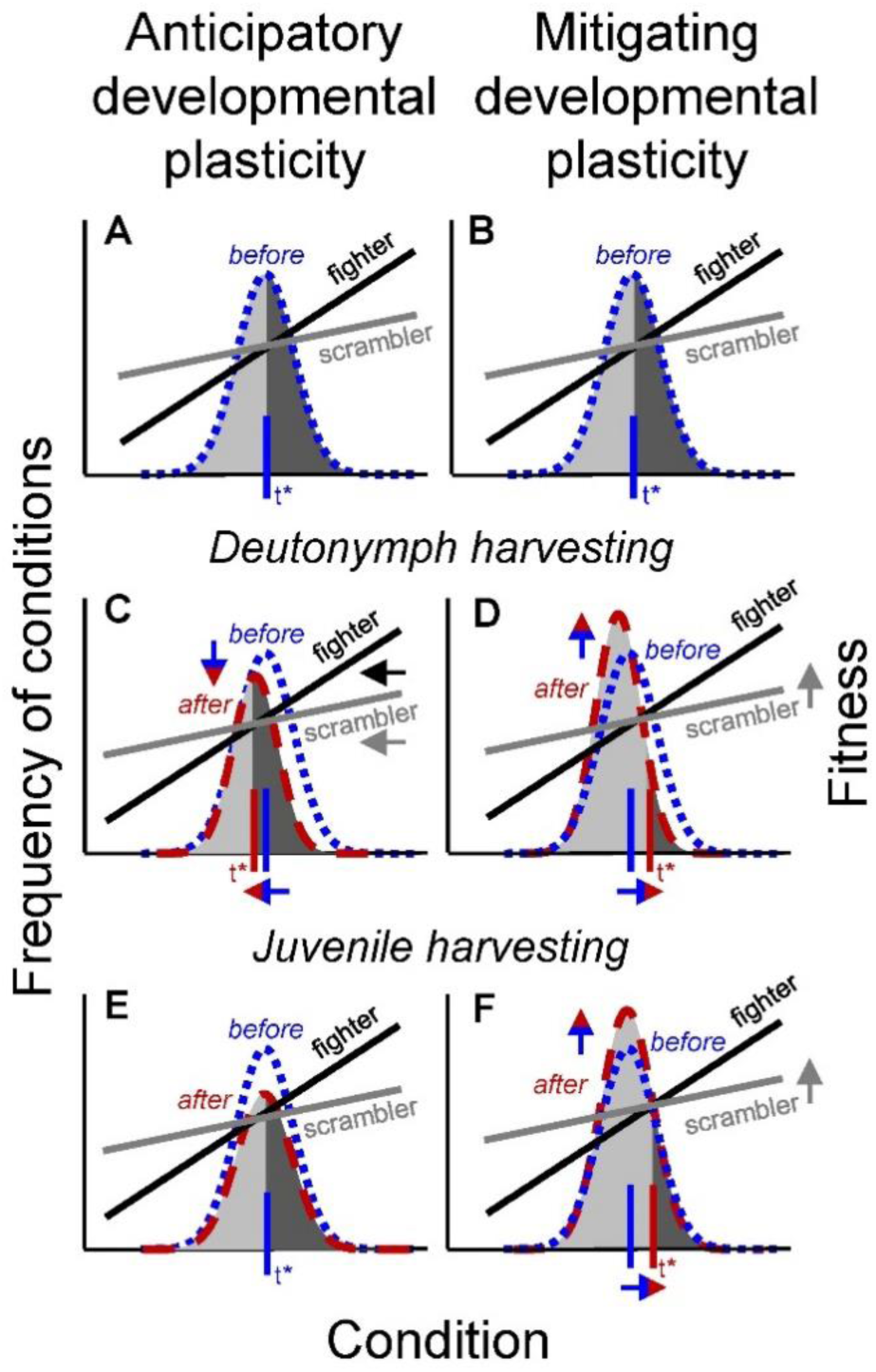
Eco-evolutionary changes due to the selective harvesting of deutonymphs and other juveniles. According to the ET model (Hazel et al. 1990; 2004), fitness functions of fighters (black lines) and scramblers (grey lines) cross over a condition gradient; the area under each condition distribution curve represents population size. The proportion of individuals adopting alternative phenotypes (contrastingly shaded areas under the curves: light grey for scramblers and dark grey for fighters) is determined by the threshold (t*) condition value at the intersection of the fitness functions (indicated by a vertical line). Blue dotted lines and red dashed lines represent condition distributions before (**a,b**) and after several generations of harvesting (**c-f**), respectively. **(c)** Although the selective harvesting of deutonymphs will reduce the size of the condition distribution (i.e., population size), anticipatory developmental plasticity predicts that the fitness functions, and, thus, the threshold (t*) for fighter expression, will track the change in the condition distribution and remain in the same, position relative to the condition distribution. In this example, the threshold tracks the mean of the condition distribution. (**e**) Assuming that the harvesting of juveniles reduces population size but does not change mean condition, the threshold (t*) for fighter expression remains unaffected in case of anticipatory developmental plasticity. (**d,f**) In contrast, the stress mechanism of a developmental system can facilitate the mitigating response to increased deutonymph or juvenile mortality, because males can mature early as a minor to escape the juvenile stage quickly. This will increase scrambler fitness, fuelling the evolution of scrambler expression to a higher threshold (t*) for fighter expression. Additionally, because scramblers mature earlier in life (Smallegange 2011), live longer than fighters (Radwan & Bogacz 2000) and sire more offspring than fighters (van den Beuken et al. 2019), an increase in scrambler expression can increase population size (Smallegange & Deere 2014), and thus increase the size of the condition distribution.

In contrast, according to the ET model and assuming that the male polyphenism is mitigating, the stress mechanism of the developmental system can fuel a mitigating response to increased deutonymph and juvenile mortality because males can mature early as a minor to escape the juvenile stage quickly (Ernande et al. 2004), fuelling the evolution of the threshold for scrambler expression. If scrambler fitness increases relative to that of fighters, the threshold for fighter expression will evolve to increase, both in response to deutonymph harvesting (compare the proportions of the differently shaded areas under the condition distribution in Fig. 1b [before harvesting] with Fig. 1d [after harvesting for several generations]) and in response to the selective harvesting of other juveniles (compare the proportions of the differently shaded areas under the condition distribution in Fig. 1b [before harvesting] with Fig. 1f [after harvesting for several generations]). Further, because scramblers mature earlier in life (Smallegange 2011), live longer than fighters (Radwan & Bogacz 2000) and sire more offspring than fighters (van den Beuken et al. 2019), an increase in scrambler expression can increase population size (Smallegange & Deere 2014). This ecological response will increase the size of the condition distribution (compare blue dotted distribution size [before harvesting] with red dashed distribution size [after harvesting for several generations] in Fig. 1d and 1f). But note that the removal of good-condition juveniles when removing deutonymphs will result in the right-hand side of the distribution to be at lower condition values (compare blue dotted distribution size [before harvesting] with red dashed distribution size [after harvesting for several generations] in Fig. 1d).

## Methods

### Study system

The bulb mite has six life stages: egg, larva, protonymph, deutonymph, tritonymph and adult (Fig. 2). The deutonymph stage is a facultative, non-feeding dispersal stage and mites only develop into this stage during unfavourable conditions (i.e. low food quality and quantity) (Díaz et al 2000). When growing from one stage to the next, mites moult; this stage is known as the quiescent stage which is immobile and non-feeding. Source populations were established from mites collected in 2010 from flower bulbs in storage rooms in North Holland. In our source populations, mites rarely develop into a deutonymph and we previously estimated the percentage of males developing into one after removing food and imposing a dry period at 3% (Smallegange & Coulson 2011). Mites were kept on an oats diet and maintained as described in Smallegange (2011).

**Figure 2.**
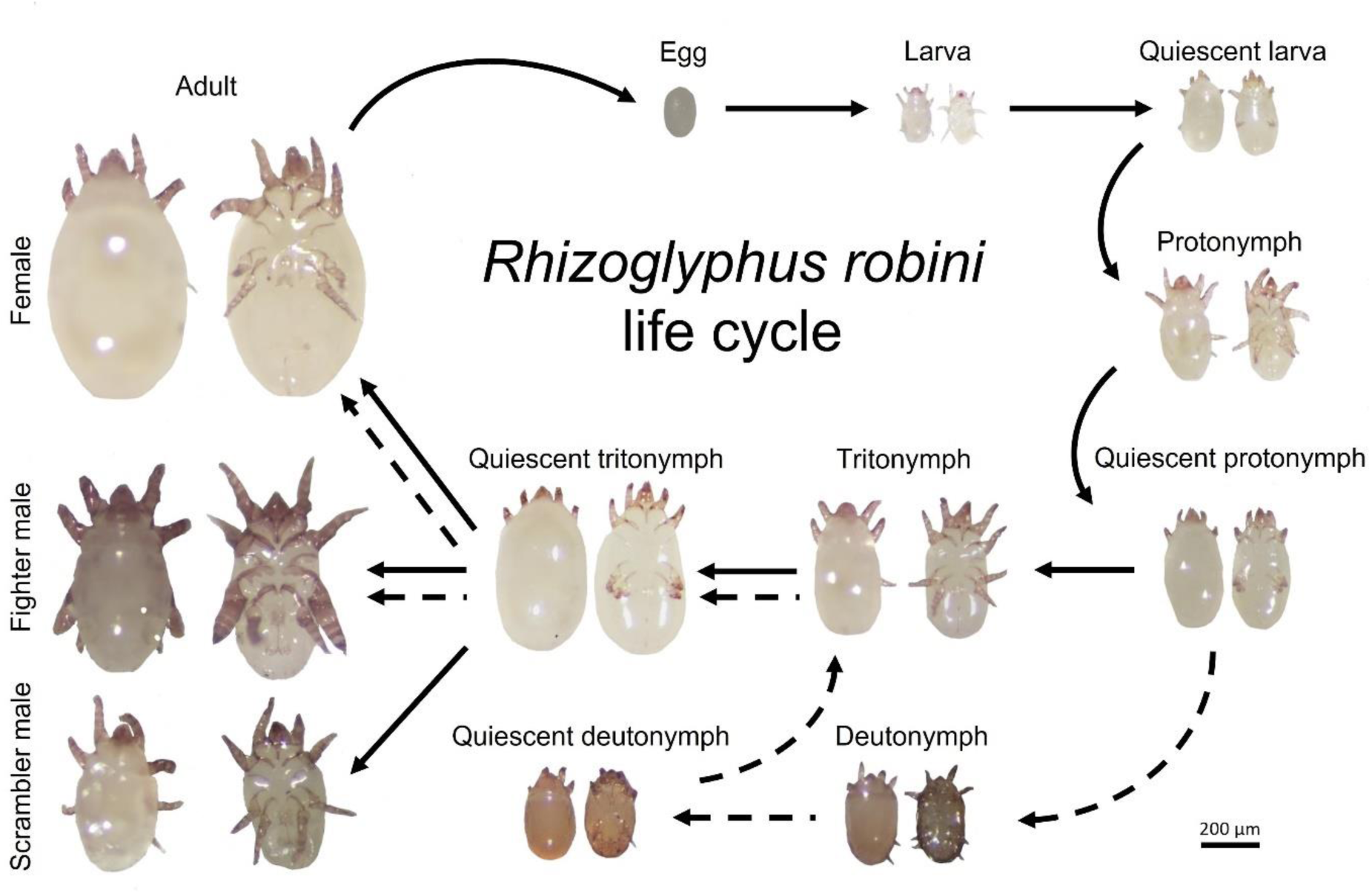
Life cycle of the bulb mite *Rhizoglyphus robini*. The bulb mite has six life stages: egg, larva, protonymph, deutonymph, tritonymph and adult. The deutonymph stage is a facultative, non-feeding dispersal stage and mites only develop into this stage during unfavourable conditions (i.e., low food quality and quantity) (Díaz et al 2000). When growing from one stage to the next, mites moult; this stage is known as the quiescent stage which is immobile and non-feeding. Show are ventral and dorsal images for each stage.

Experimental treatments were established from these source populations. Individuals across populations are highly inbred except for scrambler males in food-limited environments (Stewart et al. 2019). Inbred fighter males arise due to a combination of mating monopolization and increased survival, compared to scrambler males, which limit the genetic pool in the resulting fighter offspring (Stewart et al. 2019). The driver for inbreeding in females is likely driven by intralocus sexual conflict given the mating monopolization of fighter males (Stewart et al. 2019).

### Experimental procedure

The experiment ran between June 2016 and April 2017 (302 days) and comprised five treatments (Fig. 3): weekly harvesting of (i) 100% of all deutonymphs (D100), (ii) 50% of deutonymphs (D50), (iii) other juveniles (excluding eggs and deutonymphs) at the same percentage at which deutonymphs were harvested from all juveniles in D100 (J-D100) to keep the demographic impact of harvesting comparable over the length of the experiment, (iv) other juveniles (excluding eggs and deutonymphs) at the same percentage at which deutonymphs were harvested from all juveniles in D50 (J-D50), again, to keep the demographic impact of harvesting comparable over the length of the experiment, and (v) no harvesting (C; general control treatment). For the removal of ‘other juveniles’ we first calculated the average number of deutonymphs removed, across all three replicates, in a deutonymph removal treatment (D100/D50). Second, we calculated the average number of protonymphs and tritonymphs (i.e., other juveniles), across all three replicates, in the same deutonymph removal treatment. From these values we calculated a deutonymph proportion value: proportion value = average deutonymphs/(average protonymphs + average tritonymphs). Finally, for a treatment where other juveniles were removed (J-D100/J-D50), we multiplied the deutonymph proportion value calculated for a deutonymph removal treatment (D100/D50) to the sum of the protonymph and tritonymphs within each replicate. In the case of the ‘other juveniles’ treatments, juveniles (i.e., protonymphs and tritonymphs) were removed randomly (see further Fig. S6 in the online appendix). Because deutonymph numbers are generally low in our source populations (Smallegange & Coulson 2011), one treatment was to remove all deutonymphs. Two levels of harvesting deutonymphs (D100, D50) and juveniles (J-D100 and J-D50) were chosen to be able to compare harvesting impacts. Because we expect deutonymph numbers to be low, this means that in the other juvenile harvesting treatments, we expect to harvest low numbers as well. We anticipate that these harvesting regimes will still induce population responses. For example, in previous experiments, we also imposed harvesting regimes within which we removed only a few individuals on a regular basis and found these selections to have significant impacts on the threshold for male adult phenotype expression and population size-structure (Smallegange & Deere 2014; Smallegange & Ens 2018). Treatments were replicated three times resulting in fifteen populations. The juvenile harvesting populations were started 8 weeks after the other treatments, but before harvesting commenced (Fig. 4), for logistical reasons. At the point where the juvenile harvesting populations matched the total population sizes of the other treatments (Fig. 4), harvesting started. Populations were kept in 20 mm diameter, flat-bottomed glass tubes with a plaster of Paris and powdered charcoal base, which was kept moist to avoid desiccation of the mites. Tubes were sealed by a circle of very fine mesh (allowing gaseous diffusion), which was held in place by the tubes’ standard plastic caps with ventilation holes cut into them. Populations were kept in an unlit climate chamber at 25°C and >70% relative humidity.

**Figure 3.**
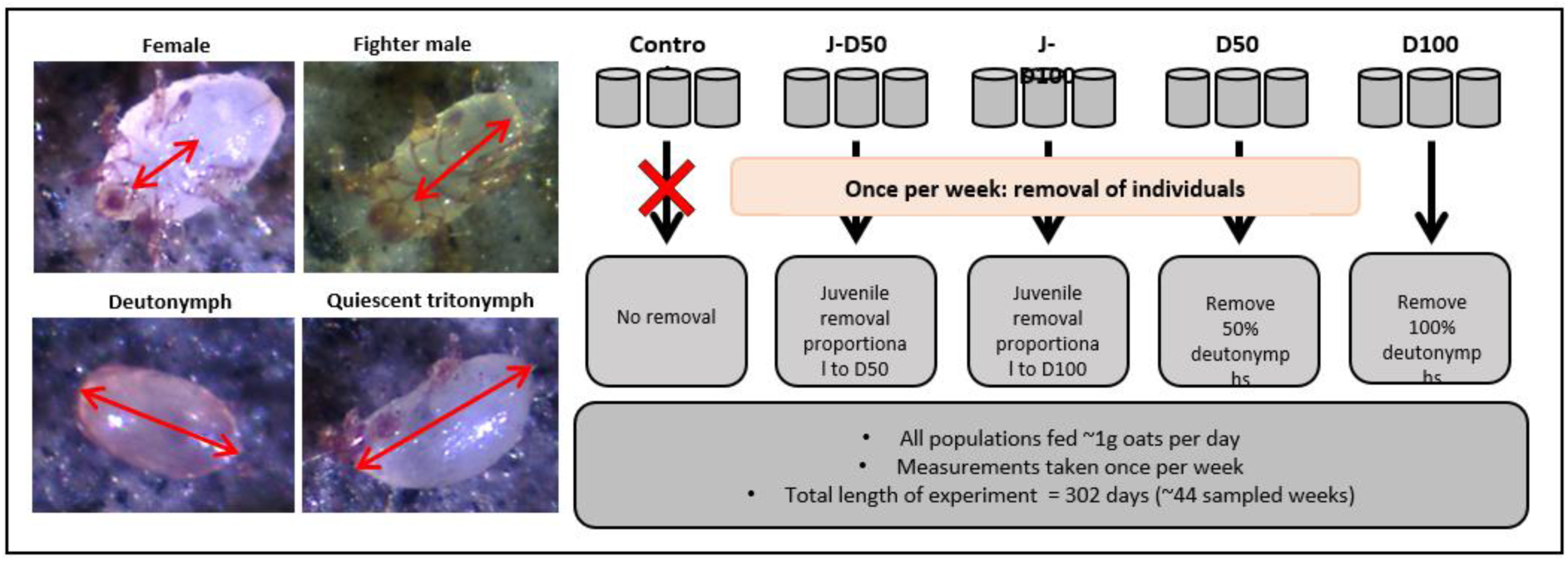
Experimental design. The photos show the length measurements of adult females (top left), adult males (top right), deutonymphs (bottom left) and quiescent individuals (bottom right). The schematic shows the experimental design and harvesting treatments.

**Figure 4.**
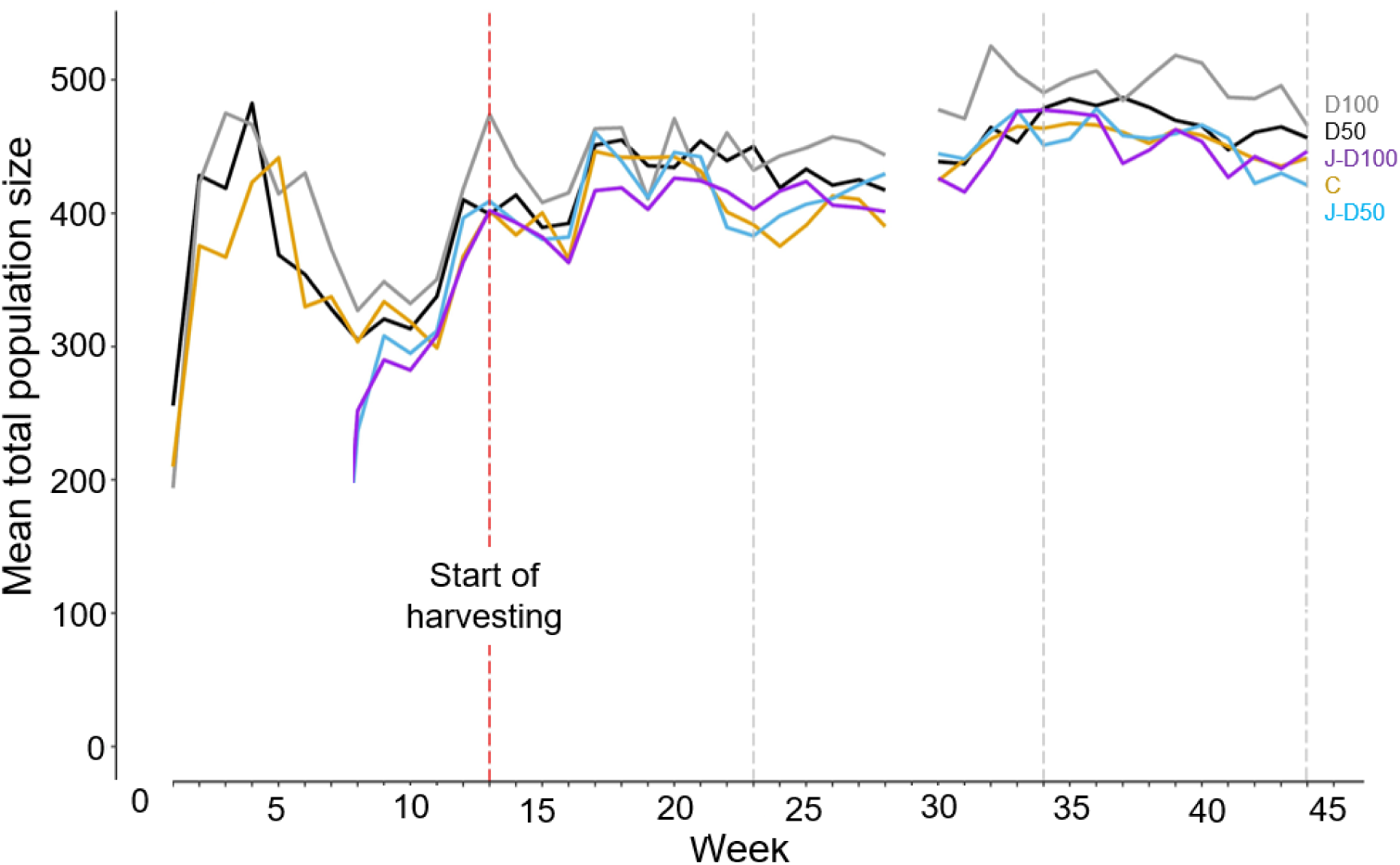
Time series of mean total population size, averaged per harvesting treatment. The treatments were: harvesting of 100% of all deutonymphs (D100: grey line), harvesting of 50% of deutonymphs (D50: black line), harvesting of other juveniles (excluding deutonymphs) at the same percentage at which deutonymphs where harvested from all juveniles in D100 (J-D100: purple line), harvesting of other juveniles (excluding deutonymphs) at the same percentage at which deutonymphs where harvested from all juveniles in D50 (J-D50: indigo line), and no harvesting (C; control treatment: orange line). The harvesting treatments began in week 13 of the experiment. No data were collected in week 29 of the experiment due to illness. Vertical grey lines denote the beginning and end of each time period over which we averaged results (see main text).

Each population was established with 50 randomly selected mites from the source population, which we refer to as founder mites, which we removed after 15 days to reduce founder effects (founder mites can be easily distinguished because they are much bigger than the first generation emerging after 15 days in the experimental population). Each population was fed approximately 1mg of oats per day (the weight was never less than 0.9g and never exceeded 1.1g). After population initialization, on each Thursday, mites of each life stage in all population were counted using a hand counter at 15 X magnification. For three weeks at the start of each time period (day 85-161, day 162-238 and day 239-302: see statistical analyses below), photos were taken of up to 5 (average: 4.05 ± 1.46 SD) individual deutonymphs, fighters, scramblers, and females and their sizes measured as proxies of condition (Rhebergen et al. 2022) to assess any shifts in mean condition across the different life stages in response to experimental conditions. The body length of mites on each photo was measured to the nearest 0.1 μm using ZEN software blue edition (ZEISS microscopy, Carl Zeiss, Germany). We measured body length for adult mites as the ventral distance of the idiosoma (the white bulbous posterior part of the body; from the mouthparts to the furthest part of the genitalia); for deutonymphs as the dorsal anterior-posterior length (without mouthparts); and for quiescent tritonymphs as the lateral body length (without mouthparts) (Fig. 3). Harvesting treatments started 85 days after populations were initialized, which is a sufficiently long time for population numbers to stabilise (which typically happens at 40 days after initialisation [Cameron and Benton 2004; Smallegange and Deere 2014]) (see also Fig. 1). Harvesting was done once a week (on Thursday), after counting and before feeding. The number of individuals harvested in the control populations (J-D100 and J-D50) were calculated based on the proportion of deutonymphs removed from the D100 and D50 populations respectively. The proportion of deutonymphs removed was calculated relative to the combined number of protonymphs and tritonymphs. This proportion value was then applied to the combined number of protonymphs and tritonymphs in the relevant control populations (proportion value for D100 (D50) was applied to J-D100 (J-D50)). The proportion value applied to the populations calculated the total number of individuals to remove from the J-D100 and J-D50 populations. Only protonymphs and tritonymphs were removed from the control populations and this was done randomly. The total protonymphs and deutonymphs removed for each population for each week can be found in data file in the online appendix (Smallegange and Deere 2023). The harvesting treatments were applied for a total of 217 days, which in a similar experiment was sufficient to observe an evolutionary shift in fighter expression (Smallegange & Deere 2014).

However, even if this period of time is too short to observe an evolutionary response to harvesting, we can still distinguish between our anticipatory and mitigating hypotheses. Specifically, under juvenile harvesting (J-D100 and J-D50), fighter expression is predicted to remain unaltered if it is anticipatory (Fig. 1e). However, fighter expression is predicted to decrease if it is mitigating (Fig. 1f) because juvenile males can respond quickly (within a generation) to increased juvenile mortality by expediating ontogeny and mature as a scrambler. This plastic (ecological) response, in turn, can further fuel the evolution towards developmental systems that produce scramblers in response to the juvenile harvesting selection pressure. What is important to note is that the ET model states that any change in the threshold for alternative male phenotype expression will affect the proportion of individuals developing either phenotype because it is expected to track the intersection of the alternative phenotype fitness functions. Therefore, we can interpret evolutionary shifts in the threshold from evolutionary changes in fighter expression (the proportion of adult males that are fighters).

### Life-history assays

Following the end of the long-term population experiment we conducted a common garden life history assay to assess if any differences in body size (which we assume to approximate condition) and fighter expression between treatments were plastic or genetic. Logistical constraints prevented us from conducting the time-consuming and labour-intensive assays to measure the threshold for alternative male phenotype expression itself. For each assay we removed ten adult females from each population, individually isolated them and gave them ad lib access to oats and allowed them to lay eggs. Three of the ten females (isolating more individuals was logistically not possible) were then randomly selected and their offspring individually isolated, given ad lib access to oats, and followed until they reached maturity. Deutonymphs, quiescent tritonymphs and adults (2-5 days after maturation) were size-measured as above and their sex and morph scored (we measured a total of 420 individuals). The effects of maternal nutritional conditions on size, age and adult male phenotype are negligible in this species (the effect size of offspring environment is 15 times larger than that of maternal environment: Smallegange 2011a), therefore, rearing mites in a common garden environment condition for one generation is sufficient to eliminate maternal effects. Females and their offspring were kept in 10 mm diameter tubes with a plaster of Paris and powdered charcoal base, which were kept in an unlit climate chamber at 25°C and >70% relative humidity.

### Statistical analyses

To analyse the results from the population experiment, we only used data from day 85 onwards (when harvesting commenced) and excluded two population tubes that had fallen over during the population experiment (one tube in treatment D100 and one in treatment J-D50). We then divided the experimental period into three periods, day 85-161, day 162-238 and day 239-302 to assess long-term (instead of transient) temporal changes in the response variables within each treatment group. To analyse the results of the population experiment, we used a generalised linear mixed model (GLMM) with Gaussian errors to analyse the effect of the fixed factors harvesting treatment (C, D100, D50, J-D100, J-D50) and time period (period 1, 2, and 3) and their interaction, including ‘population tube’ as a random effect (to account for the repeated measures within each experimental population) on: fighter expression (the number of fighters with the number of adult males set as an offset so that the model prediction will be proportion of fighter males), and the mean size of deutonymphs, adult fighters, scramblers and females. The model assumptions of Gaussian errors and homoscedacity were confirmed by inspecting the probability plots and error structures. We used a GLMM with Poisson errors to analyse the effects of the fixed factors harvesting treatment and time period, their interaction, with ‘population tube’ as a random effect, on the number of deutonymphs and on total population size (all individuals). In the online appendix (Smallegange and Deere 2023) we present the analysis of treatment effects on the number of juveniles and female and male adults.

To analyse the results of the life history assay conducted at the end of the population experiment, we used a GLMM with Gaussian errors to analyse the effect of the fixed factor harvesting treatment (C, D100, D50, C-D100, C-D50), including ‘population tube’ and ‘maternal identity’ as random effects on the mean size of adult fighters, scramblers and females. The model assumptions of Gaussian errors and homoscedacity were again confirmed by inspecting the probability plots and error structures. We used a GLMM with binomial errors to analyse the effect of the harvesting treatment on fighter expression (‘0’ if a male developed into a scrambler, and ‘1’ if in a fighter), including ‘population tube’ and ‘maternal identity’ as random effects.

In each GLMM, the significance of the treatment effects was tested using a model simplification procedure, whereby the full model was fitted after the least significant term had been removed (never deleting random terms from the model), if the deletion caused an insignificant increase in deviance as assessed by a maximum likelihood ratio test (LRT). Pairwise comparisons among the treatment levels were conducted by creating a reduced model within which two levels were combined. If the reduced model caused an insignificant increase in deviance as assessed by a maximum likelihood ratio test (α = 0.05), then the two levels were considered to be not significantly different. All analyses were conducted in R and for all GLMMs we used the *lme4* package (www-r-project.org).

## Results

### Population size, fighter expression and number of deutonymphs

Mean total population size slightly increased over time since we started harvesting (Fig. 4).

Indeed, mean population size significantly differed between time periods (χ_2_^2^= 262.07, p<0.001) (non-significant interaction: χ_8_^2^= 7.30, p=0.063) and was also significantly affected by harvesting (χ_4_^2^ = 15.58, p=0.004). Population size was highest when 100% of deutonymphs were harvested (D100), lower when 50% of deutonymphs were harvested (D50), and lowest in all the other treatments (C, J-D50 and J-D100) (treatments C and J-D50 did not differ significantly (χ_1_^2^ = 0.35, p=0.553), and neither did C and J-D50 differ significantly from J-D100 (χ_1_^2^= 0.47, p=0.494)) (Fig. 5A). Irrespective of harvesting treatment, total population size significantly increased with increasing time period (Fig. 5B). The harvesting treatment significantly affected the proportion of fighters but differently at different time periods (significant interaction harvesting × time period: χ_8_^2^ = 29.92, p<0.001). Within the interaction, the harvesting treatments C and J-D50 did not significantly differ in their effect on proportion of fighters in the different time periods (χ^2^= 4.22, p=0.239) and were combined into one level. The next two similar responses to the harvesting treatments J-D100 and D50 did, however, differ significantly (χ_3_^2^= 11.80, p=0.008) (Fig. 5C). In summary, the proportion of fighters was always relatively the lowest when 100% of deutonymphs were harvested (D100 treatment) (Fig. 5C), but when other juveniles were harvested at the same percentage as in D100, (J-D100), and when harvesting 50% of deutonymphs (D50), the proportion of fighters was relatively high in the first time periods, but lower in the last time period (Fig. 5C). In contrast, under the combined harvesting treatments of C and J-D50, the proportion of fighters was always high over the whole course of the experiment (Fig. 5C). In the online appendix we present the number of fighters and scramblers for each harvesting treatment (Smallegange and Deere 2023).

**Figure 5.**
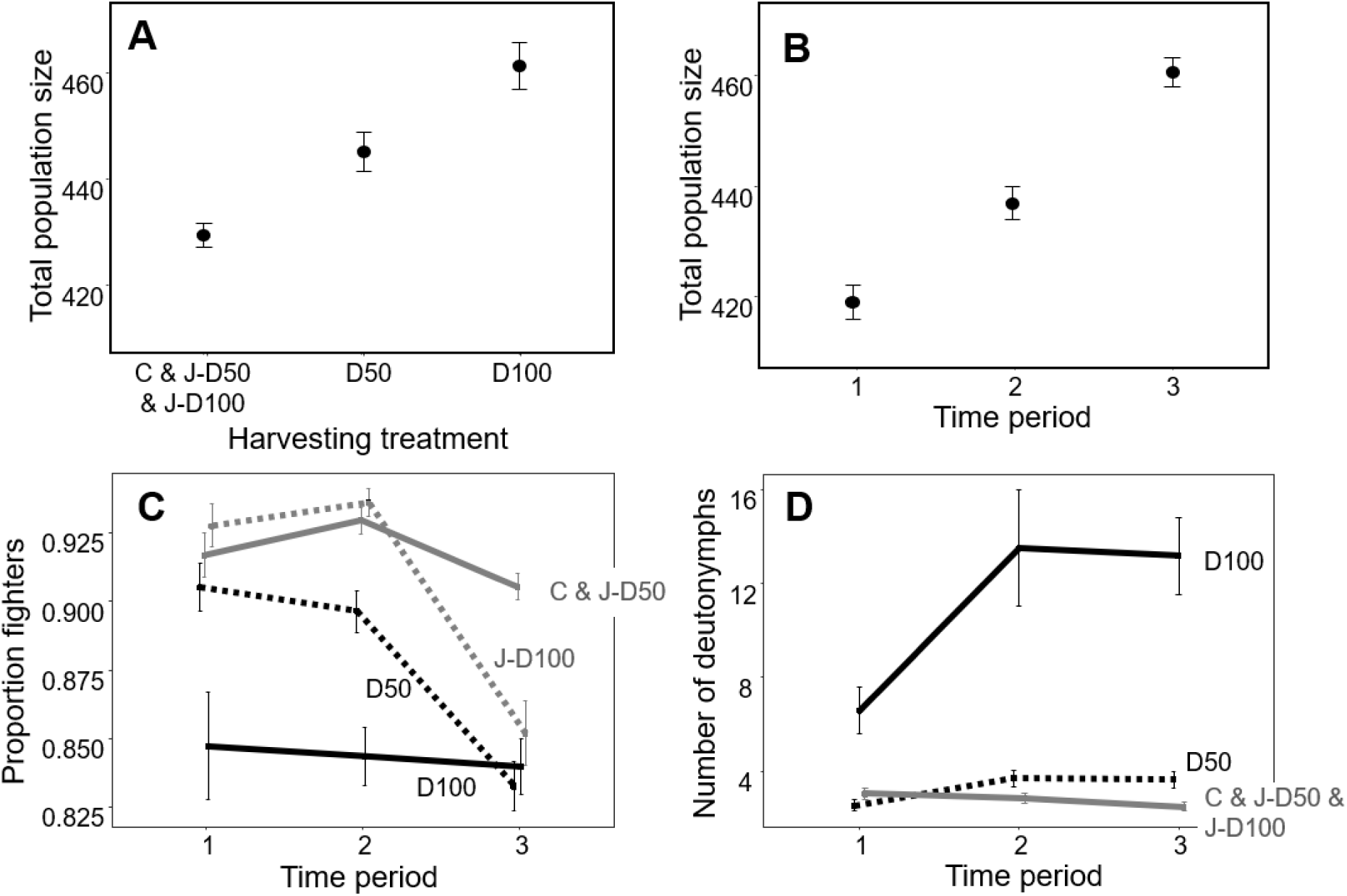
Population size and structure of the long-term population experiment. Mean total population size differed between different harvesting treatments **(a)** and between the different time periods **(b)**. Fighter expression varied over time in most but not all harvesting treatments **(c)**. The number of deutonymphs also varied over time in some but not all harvesting treatments **(d)**. Errors in all panels are standard errors. Harvesting treatments were: harvesting of 100% of all deutonymphs (D100: solid black lines), harvesting of 50% of deutonymphs (D50: dashed black lines), harvesting of other juveniles (excluding deutonymphs) at the same percentage at which deutonymphs where harvested from all juveniles in D100 (J-D100: grey dashed line (c) and solid grey line (d)), harvesting of other juveniles (excluding deutonymphs) at the same percentage at which deutonymphs where harvested from all juveniles in D50 (J-D50: solid grey lines), and no harvesting (C; control treatment: solid grey lines). Treatment levels that did not significantly differ from each other are grouped (see main text). Note that y-axes do not start at zero.

The harvesting treatment also significantly affected the number of deutonymphs, but, again, differently at different time periods (significant interaction harvesting × time period: χ_8_^2^ = 49.70, p<0.001). Within the interaction, the harvesting treatment levels J-D100 and C did not significantly differ (χ_3_^2^= 0.40, p=0.940) and were combined into one level. Likewise, the latter combined level did not significantly differ from the J-D50 harvesting treatment (χ_3_^2^ = 3.08, p=0.380) and all three control levels were combined into one level. These three levels, combined, significantly differed from the D50 harvesting level (χ_3_^2^ = 12.42, p=0.006) and from the D100 harvesting level (χ_3_^2^ = 67.71, p<0.001). We therefore find that, counterintuitively, the number of deutonymphs was higher in the last time period than in the first one when all deutonymphs were harvested (D100), and, to a lesser extent, when 50% of deutonymphs were harvested (D50) (Fig. 5D). In contrast, in the other treatments combined, the number of deutonymphs was slightly lower in the last time period than in the first (Fig. 5D). Furthermore, at any given time period, the number of deutonymphs was always the highest in the treatment where weekly harvesting removed all deutonymphs (D100) (Fig. 5D). In the online appendix (Smallegange and Deere 2023), we present interaction plots for the three response variables.

### Deutonymph, fighter, scrambler and adult female mean size

Mean deutonymph size was not significantly affected by harvesting (χ_4_^2^ = 4.81, p=0.307) but decreased with each consecutive time period (significant effect of time period: χ_2_^2^ = 106.31, p<0.001) (Fig. 6A) (non-significant interaction: χ_8_^2^ = 7.22, p=0.513). Mean fighter size showed a similar response but size differed significantly between the different harvesting treatments (significant interaction harvesting × time period: χ_8_^2^ = 19.44, p=0.013) (Fig. 6B). Within the interaction, fighter size in the harvesting treatment levels J-D50 and J-D100 did not significantly differ (χ_3_^2^= 2.33, p=0.507) and were combined into one level (Fig. 6B). Similarly, fighter size in the harvesting treatment levels C and D50 did not significantly differ (χ_3_^2^= 1.83, p=0.609) and were combined into one level (Fig. 6B). The overall result was that mean fighter size in the combined levels J-D50 and J-D100 and in the combined levels C and D50 was lower in the last than in the first time period, whereas mean fighter size in the D100 treatment did not decrease until the last time period (Fig. 6B). Mean scrambler size was not significantly affected by harvesting (χ_4_^2^ = 1.17, p=0.883) but was significantly lower in each consecutive time period (significant effect of time period: χ_2_^2^= 96.44, p<0.001) (Fig. 6C) (non-significant interaction: χ_8_^2^ = 9.48, p=0.303). Finally, mean adult female size was also not significantly affected by harvesting (χ_4_^2^ = 0.62, p=0.961) but significantly decreased with each consecutive time period (significant effect of time period: χ_2_^2^= 67.65, p<0.001) (Fig. 6D) (non-significant interaction: χ_8_^2^ = 14.55, p=0.068). In the online appendix (Smallegange and Deere 2023), we present interaction plots for the four response variables.

**Figure 6.**
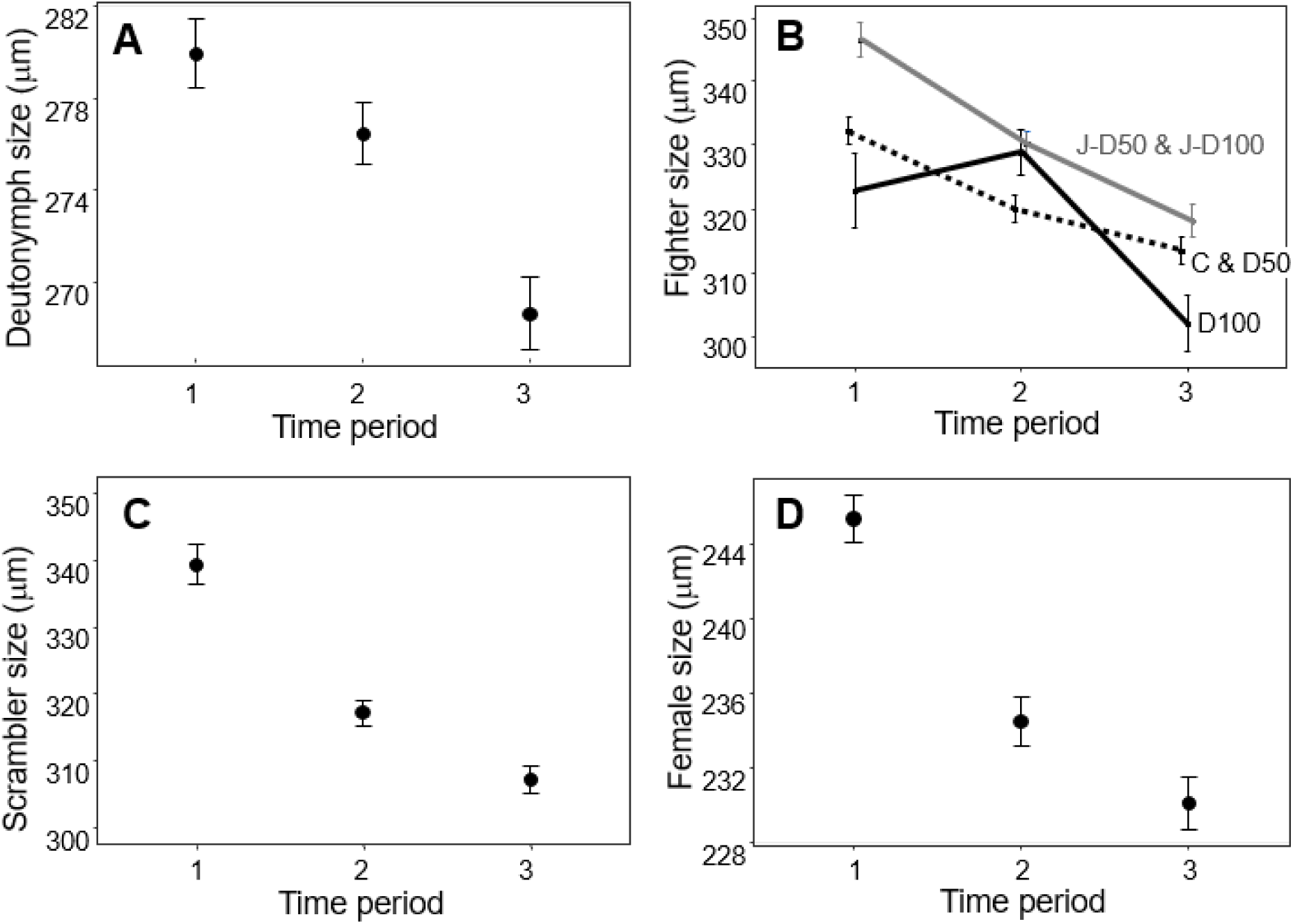
Mean body sizes observed in the long-term population experiment. Mean deutonymph size (μm) **(a)**, mean fighter size (μm) **(b)**, mean scrambler size (μm) **(c)** and mean female size (μm) **(d)** all significantly decreased over the course of the experiment, but for fighters, this decrease differed between harvesting treatments (d). Errors in all panels are standard errors. Harvesting treatments were: harvesting of 100% of all deutonymphs (D100: solid black line), harvesting of 50% of deutonymphs (D50: dashed black line), harvesting of other juveniles (excluding deutonymphs) at the same percentage at which deutonymphs where harvested from all juveniles in D100 (J-D100: grey solid line), harvesting of other juveniles (excluding deutonymphs) at the same percentage at which deutonymphs where harvested from all juveniles in D50 (J-D50: grey solid line), and no harvesting (C; control treatment: dashed black line). Treatment levels that did not significantly differ from each other are grouped (see main text). Note that y-axes do not start at zero.

### Life history assay

Neither fighter size (χ_4_^2^ = 6.58, p=0.160, mean size: 439.76 μm ± 2.39 SE, n = 190) (Fig. 7A), scrambler size (χ_4_^2^ = 2.77, p=0.597, mean size: 401.07 μm ± 7.33 SE, n = 14) (Fig. 7B) or female adult size (χ_4_^2^= 3.75, p=0.440, mean size: 334.66 μm ± 1.65 SE, n = 206) (Fig. 7C) were affected by the harvesting treatments in the common garden life history assay. Fighter expression was also not significantly affected by harvesting treatment (χ^2^ = 0.51, p=0.973) (Fig. 7D); however, only 14 scramblers emerged in the life history assay (as opposed to 190 fighters), prohibiting any statistical inference.

**Figure 7.**
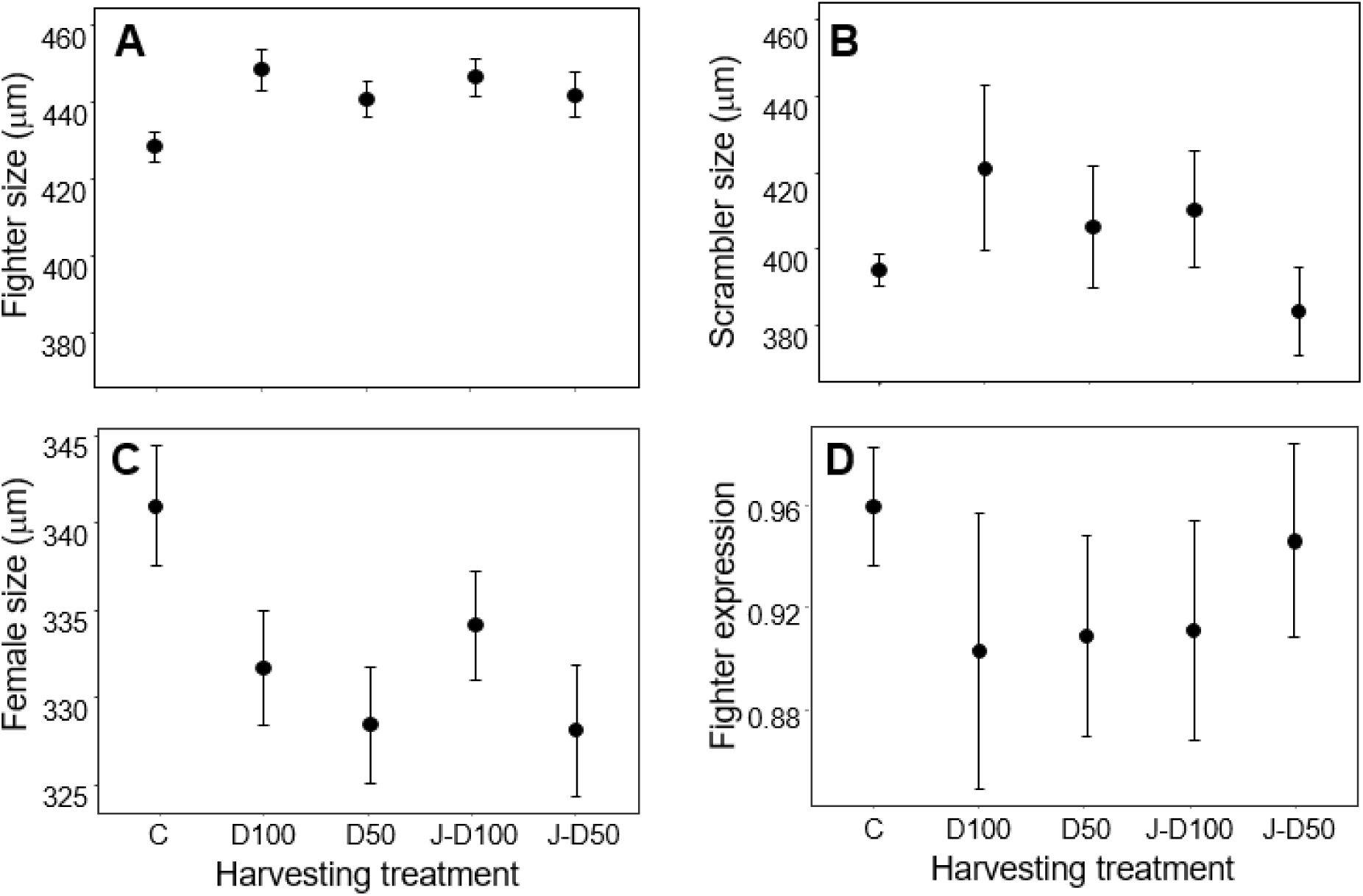
Life history assay results. Mean fighter size (μm) (**a**), scrambler size (μm) **(b)**, female size (μm) **(c)**, and fighter expression (proportion of adult males that are fighters) **(d)** for each harvesting treatment. Errors in all panels are standard errors. Harvesting treatments were: harvesting of 100% of all deutonymphs (D100), harvesting of 50% of deutonymphs (D50), harvesting of other juveniles (excluding deutonymphs) at the same percentage at which deutonymphs where harvested from all juveniles in D100 (J-D100), harvesting of other juveniles (excluding deutonymphs) at the same percentage at which deutonymphs where harvested from all juveniles in D50 (J-D50), and no harvesting (C; control treatment). Note that y-axes do not start at zero and differ between panels.

## Discussion

We tested two hypotheses on how developmental plasticity impacts the eco-evolutionary response of male polyphenic bulb mite populations to the selective removal of individuals of different developmental stages. If alternative male phenotype expression is taken to be anticipatory (anticipating future mating success), we predicted that the selective harvesting of deutonymphs and other juveniles would not impact fighter expression. In contrast, if alternative male phenotype expression is taken to be mitigating, we predicted that the selective harvesting of deutonymphs and other juveniles would decrease fighter expression both plastically, as male juveniles can expediate development and escape the juvenile stage quicker by maturing as a scrambler, and evolutionarily, as the selective removal of juveniles fuels the evolution of developmental systems that produce scramblers. In line with the mitigating hypothesis, we found that fighter expression was overall lowest when 100% of all deutonymphs were harvested (D100), and, at the end of the experiments, had reached similarly low levels in the treatments where 50% of deutonymphs were harvested (D50) and where the highest number of other juveniles were harvested (J-D100). This ‘delayed’ response could be due to that fact that our proportional harvesting treatments became more severe as population size increased over the course of the experiment, whereas per capita food availability would also have decreased with increasing population size, potentially reducing fighter expression. In contrast, fighter expression remained high throughout the experiment in the control treatment and in the treatment where the lowest number of juveniles were harvested (J-D50), perhaps because, there, the harvesting impact was the lowest. Our life history assay at the end of the population experiment failed at showing any sign of evolutionary differentiation between our treatments, which could be due to low statistical power, no evolution taking place or insufficient initial genetic diversity. Also, if the overall observed reduction in body size (condition) affected deutonymph numbers, this plastic response could have reduced the strength of the selection imposed by our selective harvesting of deutonymphs. However, it is striking that the harvesting treatments where we observed the lowest levels of fighter expression in the population experiment are the same as where we observed the lowest fighter expression levels in the life history assay. But even if this pattern was coincidental and not indicative of an evolutionary change in fighter expression, our findings still point to a role for mitigating developmental plasticity. That is, we expected no shift in fighter expression at all in response to the harvesting of juveniles (bar deutonymphs) under anticipatory developmental plasticity (Fig. 1e), but a reduction in fighter expression under mitigating developmental plasticity (Fig. 1f) because individuals mature earlier as a scrambler to escape the risky juvenile stage. The latter reduction is indeed what we observed towards the end of the experiment in the treatment where we harvested the highest number of juveniles other than deutonymphs (J-D100). Our results therefore suggest that nutrition-deprived male juveniles do not ‘make the best of a bad job’ by anticipating a non-fighting mating tactic (Dawkins 1980; Eberhard 1982), but instead develop into that male adult phenotype because they can adaptively mitigate their early-life stress (Rhebergen 2022). Such adaptive early-life stress responses that have negative effects in adulthood are common in (warm-blooded) mammals and birds (Lindström 1999; Fisher et al. 2006; Wells 2011; Vickers 2014), likely because their growth during development is physiologically constrained. However, if we account for such pre-adult viability selection in coldblooded species that typically show flexible growth, it would allow us to better characterise natural selection on trait development like male polyphenisms in these species, how it can affect the response to other selections in adulthood (e.g., sexual selection: Mojica and Kelly 2010; Lürig & Matthews 2021; Rhebergen 2022), and how such trait dynamics influence, and are influenced by, population dynamics (Smallegange 2022).

One unexpected finding in our experiment was that the number of deutonymphs was highest when we harvested all of them, each week. Further, despite being harvested, deutonymph number increased as the experiment progressed. We can only speculate why this happened. Perhaps this overcompensatory response was because the per capita food availability increased as we removed individuals (Verhulst 1838, Schaefer 1954, Hilborn and Walters 1992), although we still only maximally removed a small percentage of the total population in the deutonymph harvesting treatments, and we did not see a similar overcompensatory response in the treatment where we harvested the same percentage of other juveniles. In fact, total population size increased as well so that per capita food intake rate would decline, cueing more individuals to develop into deutonymphs. It could also be simply a numerical response because mean total population size increased over the course of the experiment and thus there were more individuals that could develop into a deutonymph. Finally, the higher number of deutonymphs we found could also reflect eco-evolutionary dynamics in which another selection pressure, for example imposed by strong-density dependence (population size was highest in the 100% deutonymph harvesting treatment), opposed the effects of direct harvesting (Edeline & Loeuille 2021). Or, perhaps, because twice as many females emerge from deutonymphs than males (fighters) (Deere et al. 2015; Stewart & Smallegange in prep), removing deutonymphs reduces females and egg output, reducing density-dependence. Either way, failure to account for either of these eco-evolutionary processes might fundamentally hamper our ability to understand the expression of dispersal phenotypes. Another explanation is that it was the actual lack of deutonymphs and their olfactory cues that triggered deutonymph development. Olfactory cues are common in acarine mites. All acarines have a sensory site on their forelegs known as the “foretarsal sensory organ” and olfaction is one of this site’s functions (Carr & Roe 2016). If the harvesting of deutonymphs reduced the concentration of deutonymph olfactory cues, this could trigger more juveniles to develop into a deutonymph because, with fewer deutonymphs around, competition for attachment sites on insects is lower (deutonymphs disperse via phoresy on insects [Díaz et al 2000]). We realise, however, that this is speculation and olfactory cues produced by deutonymphs have yet to be tested in *R. robini*. Finally, in our previous studies (Smallegange & Coulson 2011; Deere et al. 2015), male deutonymphs always matured as a fighter, but here, increases in deutonymph numbers did not coincide with increases in fighter frequency. Perhaps this was because twice as many females emerge from deutonymphs than males (fighters) (Deere et al. 2015; Stewart & Smallegange in prep) so that our statistical analyses were unable to pick up significant differences in the few more fighters that emerged. It does highlight, however, that we still have some way to go to unravel the intricate links between deutonymph expression, alternative male phenotype expression and sex ratio and how they together drive population dynamics.

Because a change in the distribution of a trait will almost always also change the size of the distribution of that trait, it follows that trait change is accompanied by a change in population size. If a trait is heritable, this constitutes eco-evolutionary change at the population level (Smallegange & Coulson 2013). This is especially so in the case of alternative phenotypes where we expect a shift in phenotype expression to be accompanied by a change in population size because the different phenotypes often have different demographic rates. For example, in the case of bulb mite males, scramblers mature earlier in life (Smallegange 2011), live longer than fighters (Radwan & Bogacz 2000) and sire more offspring than fighters (van den Beuken et al. 2019) so that an increase (or decrease) in scrambler (or fighter) expression will increase population size. Indeed, we found the highest population size in the harvesting treatment with the lowest level of fighter expression (D100: 100% deutonymph harvesting). We suspect that the increase in population size that we observed over the course of the experiment is associated with the observed decrease in mean body size over time in response to the strong, density-dependent food conditions, because smaller organisms typically reproduce faster and can reach higher equilibrium population sizes than larger ones (e.g., Huryn & Benke 2007). Indeed, Gilbert et al. (2022) recently showed in an experiment with the protist *Tetrahymena pyriformis* that mean body size decreased in response to increased density-dependence and that this rapid change in mean body size had a larger effect on population density than the reverse. Reductions in mean body size have been observed across a range of taxa in response to adverse environmental change (Gardner et al. 2011; Sheridan & Bickford 2011). Often, these reductions are associated with increased mortality rates (e.g., in red knots *Calidris canutus* [van Gils et al., 2016]). What is more, when populations not only suffer from directional environmental change but also from selective harvesting, the removal of good condition individuals (of large size) can lead to population extinction (Knell & Martínez-Ruiz 2017). Thus, unravelling the intertwined ecological and evolutionary dynamics of population size, body size, condition and condition-dependent (alternative) phenotypes is integral to understanding eco-evolutionary population responses to change (Edeline & Loeuille 2021). If our experiment would form the basis for such further studies, we suggest using outbred populations and increasing sample size in the life history assays to be able to assess any evolutionary shifts in the actual threshold for alternative male phenotype expression, and to consider other harvesting treatments with potentially stronger selective effects, like the selective removal of a high proportion of large versus randomly selected juveniles.

## Data availability statement

The raw data, the R scripts that we used to analyse the data, and the additional analyses on population structure can be found on Figshare (Deere & Smallegange 2023).

